# metaCCA: Summary statistics-based multivariate meta-analysis of genome-wide association studies using canonical correlation analysis

**DOI:** 10.1101/022665

**Authors:** Anna Cichonska, Juho Rousu, Pekka Marttinen, Antti J Kangas, Pasi Soininen, Terho Lehtimäki, Olli T Raitakari, Marjo-Riitta Järvelin, Veikko Salomaa, Mika Ala-Korpela, Samuli Ripatti, Matti Pirinen

## Abstract

A dominant approach to genetic association studies is to perform univariate tests between genotype-phenotype pairs. However, analysing related traits together increases statistical power, and certain complex associations become detectable only when several variants are tested jointly. Currently, modest sample sizes of individual cohorts and restricted availability of individual-level genotype-phenotype data across the cohorts limit conducting multivariate tests.

We introduce *metaCCA*, a computational framework for summary statistics-based analysis of a single or multiple studies that allows multivariate representation of both genotype and phenotype. It extends the statistical technique of canonical correlation analysis to the setting where original individual-level records are not available, and employs a covariance shrinkage algorithm to achieve robustness.

Multivariate meta-analysis of two Finnish studies of nuclear magnetic resonance metabolomics by *metaCCA*, using standard univariate output from the program SNPTEST, shows an excellent agreement with the pooled individual-level analysis of original data. Motivated by strong multivariate signals in the lipid genes tested, we envision that multivariate association testing using *metaCCA* has a great potential to provide novel insights from already published summary statistics from high-throughput phenotyping technologies.

Code is available at https://github.com/aalto-ics-kepaco.

## 1 Introduction

Most human diseases and traits have a strong genetic component. Genome-wide association studies (GWAS) have proven effective in identifyin*G* genetic variation contributing to common complex dis-orders, including type 2 diabetes (Mahajan *et al.*, 2014), cardiovascular disease (Deloukas et *al.*, 2013), schizophrenia (Schizophrenia Working Group of the Psychiatric Genomics Consortium, 2014), and quantitative traits, such as lipid levels (Global Lipids Genetics Consortium, 2013; Surakka *et al.*, 2015) and metabolomics (Kettunen *et al.*, 2012; Shin *et al.*, 2014).

A dominant approach to GWAS is to test one single-nucleotide polymorphism (SNP) at a time against one quantitative phenotype measure or a binary disease indicator. This univariate approach is unlikely to be optimal when millions of SNPs and a growing number of phenotypes, including serum metabolomic profiles (Kettunen *et al.*, 2012; Shin *et al.*, 2014), three-dimensional images (Wang *et al.*, 2013), and gene expression data (Ardlie *et al.*, 2015) become available simultaneously. Indeed, a recent comparison demonstrated that utilising multivariate phenotype representation increases statistical power and leads to richer findings in the association tests compared to the univariate analysis (Inouye *et al.*, 2012). Moreover, some complex genotype-phenotype correlations can be detected only when testing several genetic variants simultaneously (Marttinen *et al.*, 2014).

Unfortunately, restricted availability of complete multivariate individual-level records across the cohorts currently limits multivariate analyses. Often, only the univariate GWAS summary statistics from individual cohorts are publicly available. Hence, a major question is how we can use the univariate results to carry out a multivariate meta-analysis of GWAS (Evangelou and Ioannidis, 2013), which is crucial to increase the power to identify novel genetic associations.

Recently, two kinds of approaches for utilising univariate summary statistics in multivariate testing have been proposed: 1) one SNP against multiple traits (Stephens, 2013; Vuckovic *et al.*, 2015; Zhu *et al.*, 2015) and 2) multiple SNPs against one trait (Vaitsiakhovich *et al.*, 2015; Feng *et al.*, 2014; Yang *et al.*, 2012). We propose a new framework, *metaCCA*, that unifies both of the existing approaches by allowing canonical correlation analysis (CCA) of multiple SNPs against multiple traits based on univariate summary statistics and publicly available databases.

CCA is a well-established statistical technique for identifying linear relationships between two sets of variables, and has been successfully applied to GWAS (Inouye *et al.*, 2012; Marttinen *et al.*, 2013; Ferreira and Purcell, 2009; Tang and Ferreira, 2012). Our *metaCCA* method extends CCA to the setting where original individual-level measurements are not available. Instead, *metaCCA* works with three pieces of the full data covariance matrix, and applies a covariance shrinkage algorithm to achieve robustness. We demonstrate the performance of *metaCCA* using SNP and metabolite data from three Finnish cohorts. In summary, this paper makes the following contributions.

- To our knowledge, we provide the first computational framework for association testing between multivariate genotype and multivariate phenotype based on univariate summary statistics from single or multiple GWAS. Our implementation is freely available.
- We demonstrate how to accurately estimate correlation structures of phenotypic and genotypic variables without an access to the individual-level data.
- We avoid false positive associations by a covariance shrinkage algorithm based on stabilisation of the leading canonical correlation.
- Our approach, *metaCCA*, is a general framework to conduct CCA when full data are not available, and therefore it is widely applicable also outside GWAS.

## 2 Methods

This section is organised as follows. First, section 2.1 explains univariate GWAS, the results of which, in the form of cross-covariance matrix, constitute an input to *metaCCA* described in section 2.2; section 2.3 demonstrates how a meta-analysis of several studies is conducted in our framework; section 2.4 outlines a procedure for choosing SNPs representative of a given locus; finally, section 2.5 introduces the data we used to test *metaCCA* in the meta-analytic setting.

### 2.1 Univariate GWAS

Let *X* and *Y* denote genotype and phenotype matrices of dimensions *N ×G* and *N ×P*, respectively, storing the individual-level data; *N* the number of samples; G and P the number of genotypic and phenotypic variables, respectively. The columns of *X and Y* are standardised to have mean 0 and standard deviation 1.

Typically, univariate GWAS analysis of quantitative traits tests for an association between each pair of genotype ***x***_*g*_ ∈ ℛ^*N*^ and phenotype ***y***_*p*_ ∈ ℛ^*N*^ separately using a linear model:

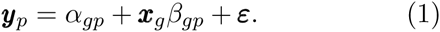

Coefficient *β*_*gp*_, corresponding to the slope of the regression line, is the parameter of interest, since it depicts the size of the effect of the genetic variant ***x***_*g*_ on the trait ***y***_p_. Parameter *α*_*gp*_ is an intercept on the y-axis, and ϵ indicates a Gaussian error term or noise. The model is fit by the method of *least squares* that leads to a closed-form estimate for the unknown parameter *β* _p_:

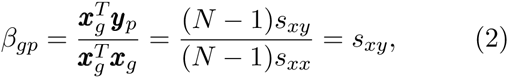

where *s*_*xy*_ is a sample covariance of ***x***_*g*_ and ***y***_*p*_, and *s*_*xx*_ = 1 is a sample variance of ***x***_*g*_. Hence, the cross covariance matrix ∑_*X Y*_ between all genotypic and phenotypic variables is made of univariate regression coefficients *β*_*gp*_:

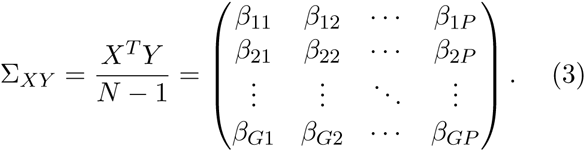

An important note is that if the individual-level data sets *X* and *Y* were not standardised before applying the linear regression, the standardisation can be achieved afterwards by a transformation

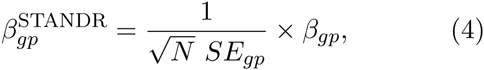

where *SE*_*gp*_ indicates the standard error of *β*_gp_, as given by GWAS software. (Typically, 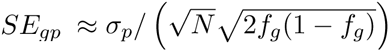, where *σ*_*p*_ is the standard deviation of the trait *p*, and *f*_*g*_ is the minor allele frequency of SNP *g*, but uncertainty in the genotype imputation causes deviations from this expression.)

### 2.2 metaCCA

Conducting multivariate association tests requires estimates of the dependencies between genotypic and phenotypic variables, denoted ∑_*X X*_ and ∑_*X Y*_, respectively. Typically, they are calculated based on the individual-level measurements *X* and *Y*:

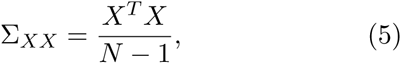

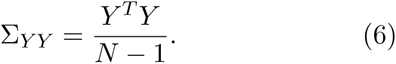

*metaCCA* operates on the cross-covariance matrix ∑_*X Y*_ (eq. 3) and correlation structures 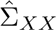, 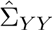, estimated without an access to the individual-level data *X* and *Y* (Figure 1A-B). To make the resulting full covariance matrix ∑ a valid covariance matrix, *metaCCA* applies a shrinkage algorithm (Figure 1C).

**Figure 1:**
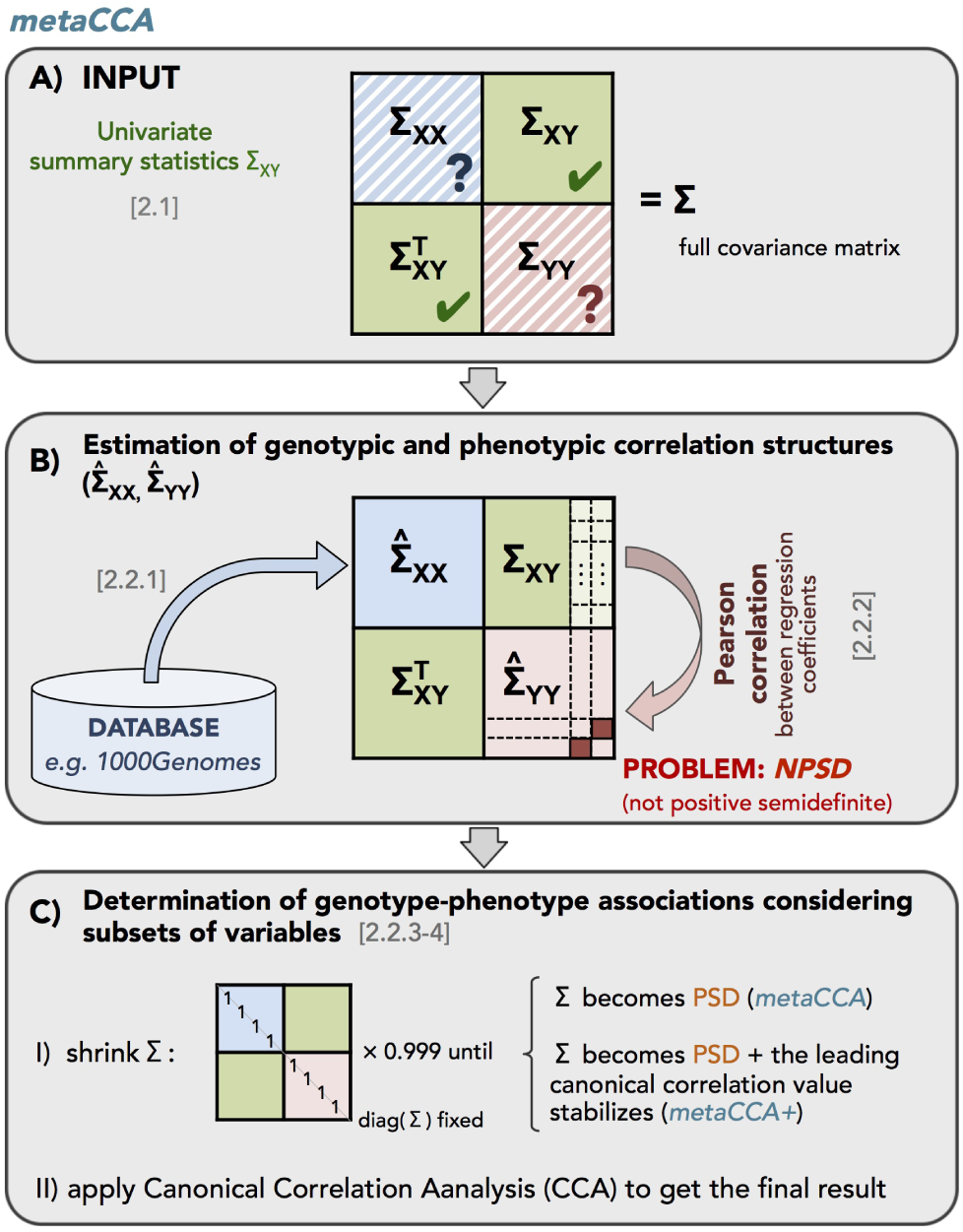
Schematic picture showing an overview of *metaCCA* framework for summary statistics-based multivariate association testing using canonical correlation analysis. A) *metaCCA* operates on three pieces of the full covariance matrix ∑: ∑ _*X Y*_ of univariate genotype-phenotype association results, ∑_*X X*_ of genotype-genotype correlations, and EYY of phenotype-phenotype correlations. B) 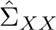 is estimated from a reference database matching the study population, e.g. the 1000 Genomes, and phenotypic correlation structure 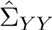 is estimated from ∑_X Y_. C) A covariance shrinkage algorithm is applied to add robustness to the method. Numbers in brackets refer to subsections in Methods. Meta-analysis of several studies is performed by pooling covariance matrices of the same type, before step C), as described in section 2.3. The data reduction achieved by *metaCCA* can be seen in Supplementary Figure 1.

The rest of this section describes the details of *metaCCA* framework.

#### 2.2.1 Estimation of genotypic correlation structure

Genetic variation is organized in haplotype blocks, whose structure is determined by mutation and recombination events, together with demographic effects, including population growth, admixture and bottlenecks (Wall and Pritchard, 2003). Hence, correlation structure of genetic variants differs between populations, such as, e.g., the Finns, Icelanders or Central Europeans. In *metaCCA*, 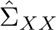 is calculated using a reference database representing the study population, such as the 1000 Genomes database (1000 Genomes Project Consortium (2012), www.1000genomes.org), or other genotypic data available on the target population. In the Results section, we demonstrate that estimating 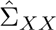 from the target population (in our case the Finns) leads to better results than utilising the data comprising individuals across distinct populations (e.g. the Finns and other Europeans). However, since reference data on the target population may not always be at hand, we also present a robust but less powerful solution to multivariate association testing by simply using genotypes of all individuals from a certain broader geographical region (e.g. a continent) available under the 1000 Genomes Project.

#### 2.2.2 Estimation of phenotypic correlation structure

In our framework, phenotypic correlation structure 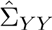 is computed based on ∑_*X Y*_. Each entry of 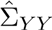 corresponds to a Pearson correlation coefficient between two column vectors of ∑_*X Y*_ - univariate summary statistics of two phenotypic variables *s* and *t* across *G* genetic variants:

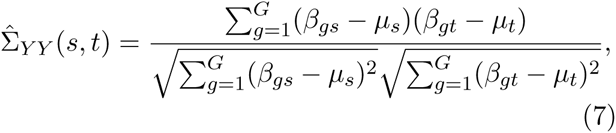

where μ_*s*_ and μ_*t*_ are the mean values computed as 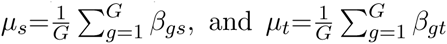. (The detailed justification is provided in Supplementary Data.) In Supplementary Data, we demonstrate that the higher the number of genotypic variables *G*, the lower the error of the estimate (Supplementary Table 2). Thus, 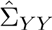 should be calculated from summary statistics for all available genetic variants, even if only a subset of them is taken to the further analysis.

#### 2.2.3 Canonical Correlation Analysis

CCA (Hotelling, 1936) is a multivariate technique for detecting linear relationships between two groups of variables *X* ∈ ℛ^*N*×*G*^ and *Y* ∈ ℛ^*N*×*P*^, where *X* and *Y* constitute two different views of the same object. The objective is to find maximally correlated linear combinations of columns of each matrix. This corresponds to finding vectors ***a*** ∈ ℛ^*G*^ and ***b*** ∈ ℛ^*P*^ that maximize

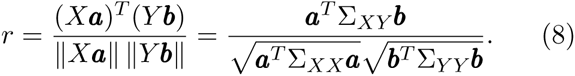

The maximised correlation *r* is called *canonical correlation* between *X* and *Y*. We provide the technical details of the method, as well as its extension to subsequent canonical correlations and their significance testing in Supplementary Data.

#### 2.2.4 Shrinkage

At this point, we have three covariance matrices, namely ∑_*X Y*_, 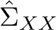and 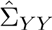. However, in most cases, the resulting full covariance matrix

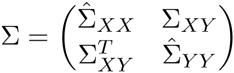

is not positive semidefinite (PSD), and therefore its building blocks cannot be just plugged into the CCA framework (eq. 8). To overcome this problem, in *metaCCA*, we apply shrinkage to find a nearest valid ∑ (Ledoit and Wolf, 2003). We use an iterative procedure where the magnitudes of the off-diagonal entries are being shrunk towards zero until ∈ becomes PSD (Algorithm 1).

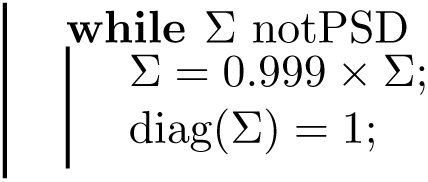
Algorithm 1

Guaranteeing the PSD property of the full covariance matrix is necessary, although, as we demonstrate in the Results section, not sufficient to obtain reliable results of the association analysis when the estimate 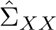(and/or 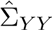) is noisy. In order to address this issue, we propose a variant of *metaCCA*, called ***metaCCA+***, where the full covariance matrix ∑ is shrunk beyond the level guaranteeing its PSD property. A challenge, however, is to find an optimal shrinkage intensity. Shrinkage applied without any stopping criterion would lead to gradual removal of all dependencies between genotypic and phenotypic variables. Ledoit and Wolf (2003) introduced an analytic approach for determining the optimal shrinkage level, but it requires the individual-level data sets *X* and *Y*. In *metaCCA+*, we monitor the leading canonical correlation value *r*, and we continue the shrinkage of the full covariance matrix ∑ until *r* stabilises. Specifically, we track the percent change *pc* of *r* between subsequent shrinkage iterations, and we determine an appropriate amount of shrinkage using an elbow heuristic, similar to the criterion for finding the number of clusters, frequently used in the literature (Tibshirani *et al.*, 2001). The idea is that the slope of the graph should be steep to the left of the elbow, but stable to the right of it. We find an elbow, and thus the appropriate number of shrinkage iterations, by taking the point closest to the origin of the plot of *pc* versus iteration number, as schematically shown in Supplementary Figure 2.

**Figure 2:**
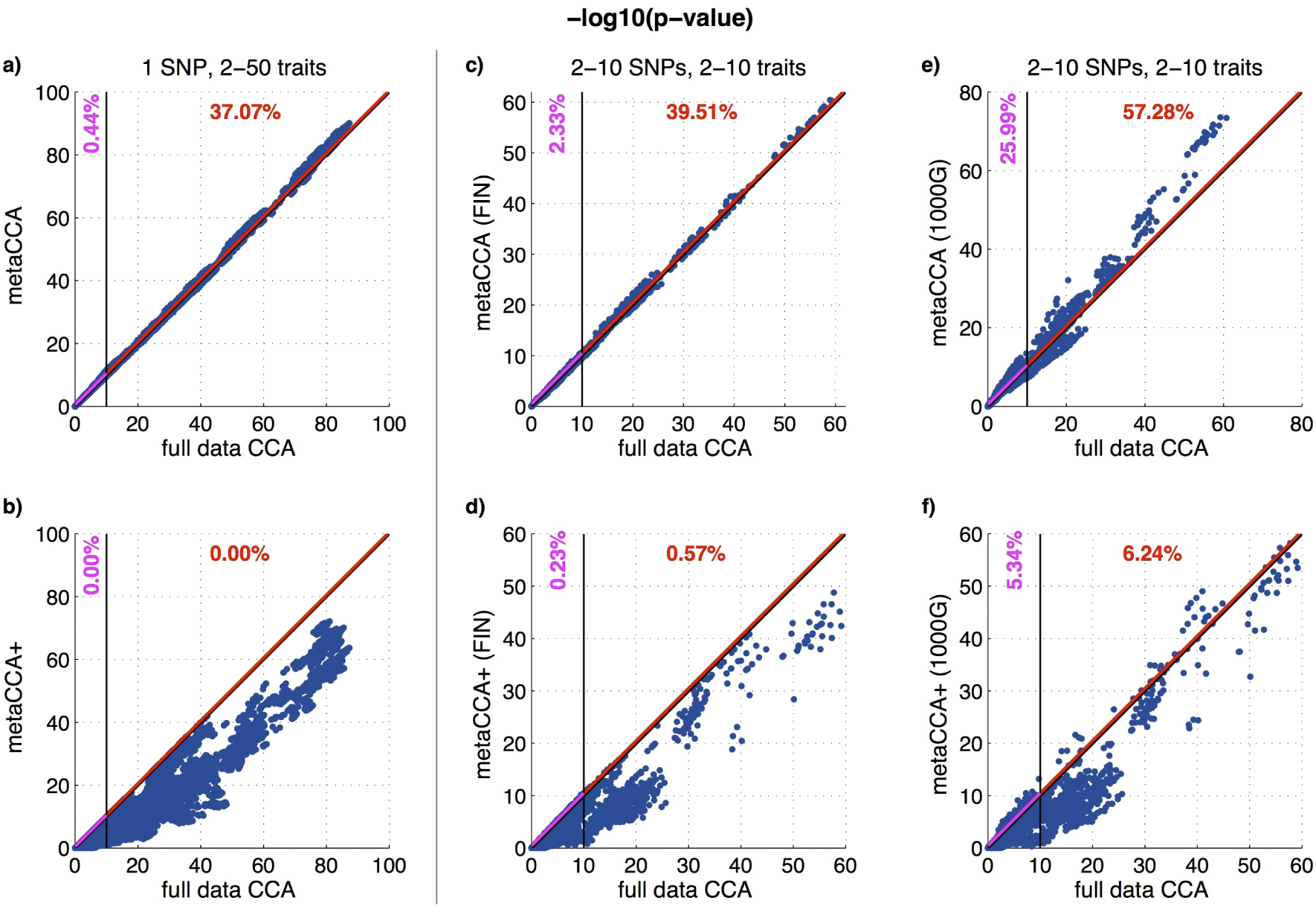
Scatter plots of −log10 p-values between the pooled individual-level analysis of original data sets *(full data CCA)* and *metaCCA* (first row), *metaCCA+* (second row). a,b) *Univariate genotype - multivariate phenotype*; meta-analysis of NFBC, FINRISK and YFS cohorts; c-f) *Multivariate genotype - multivariate phenotype;* meta-analysis of NFBC and YFS cohorts; *metaCCA/metaCCA+* was used with 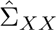 computed from FINRISK (FIN, c,d), or from the 1000 Genomes database (1000G, 503 EUR individuals, e,f). In all the cases, lipid correlation structure 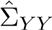 was calculated from univariate summary statistics for SNPs from the entire chromosome 1. Single point corresponds to the result of one out of a-b) 178 752, c-f) 4 050 multivariate tests. Numbers at the top of each plot indicate percentages of at least 0.5 unit overestimated *metaCCA’s/metaCCA+’s* −log10 p-values in the ranges [0, 10] (purple) or (10, max(−log10 p-value)] (red). This threshold is represented by purple and red lines. Supplementary Figure 5 shows these results restricted to the x-axis range of [0, 10].

Building blocks 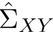, 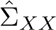, 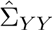 of the resulting full covariance matrix ∑, shrunk until it became PSD or beyond, are then plugged into the CCA framework to get the final genotype-phenotype association result. In the Discussion section, we further elaborate on choosing the shrinkage mode in practical applications.

#### 2.2.5 Types of the multivariate association analysis

We consider the following two types of the multivariate analysis.

1. *Univariate genotype - multivariate phenotype* One genetic variant tested for an association with a set of phenotypic variables (matrix 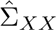 not needed).;
2. *Multivariate genotype - multivariate phenotype* A set of genetic variants tested for an association with a set of phenotypic variables.

The first type corresponds to a standard multi-trait analysis. The second type takes into account the effects across genomic variants on multiple traits, which are ignored when analysing only a single SNP or a single trait at a time.

### 2.3 Meta-analysis

*metaCCA* allows to conduct summary statistics-based multivariate analysis of one or multiple GWAS. In the meta-analytic setting, covariance matrices 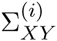, 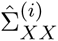 and 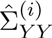 corresponding to *i*=1,…,*M* independent studies on the same topic are pooled using a weighted average:

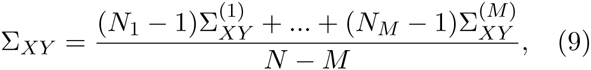

where *N*_*i*_ denotes the number of samples in the *i*^th^ cohort, and *N* = *N*_1_ +… + *N*_*M*_. This step is performed before applying the shrinkage to the full covariance matrix. As is typical for a fixed-effects meta-analysis, the weighted average is used in order to account for the varying precision of the estimates. The formulas for 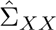 and 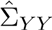 are analogous to (9). However, if all cohorts included in the meta-analysis have the same underlying population, only one genotypic correlation estimate is needed.

### 2.4 Choosing SNPs representing a locus

When analysing multiple genetic variants together, we use a procedure for selecting from a given locus a set of SNPs that jointly capture a maximal amount of genetic variation in the locus, as measured by a linkage disequilibrium (LD) score.

In each iteration, a SNP g that maximises LD-score, which we defined as *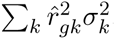*, is selected, where the sum is over all SNPs *k* that have not yet been chosen; 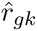 denotes a partial correlation between SNPs *g* and *k*; 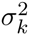 indicates empirical variance of the residuals for SNP *k* after the effects of the selected SNPs have been regressed out. The residual variance 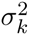 gets smaller, if the SNP has already been well explained by the previously chosen ones; hence, highly correlated SNPs will not be selected together. In the first iteration, *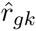* is just a Pearson correlation coefficient between SNPs *g* and *k*, and 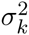 equals 1.

This procedure was used for choosing sets of SNPs representative of genes in the application of *metaCCA* to summary statistics from the program SNPTEST, described in section 3.2.

### 2.5 Data sets

In order to test our approach, we used genotypic and phenotypic data from three Finnish population cohorts: the Cardiovascular Risk in Young Finns Study (YFS, *N*_1_=2390; Raitakari *et al.* (2008)), the FINRISK study survey of 1997 (*N*_2_=3661; Vartiainen *et al.* (2010)), and the Northern Finland Birth Cohort 1966 (NFBC, *N*_*3*_=4702; Rantakallio (1969)). The detailed description of the cohorts can be found in Supplementary Data.

Our phenotype data consist of 81 lipid measures (Supplementary Table 1) from a high-throughput nuclear magnetic resonance (NMR) platform (Soininen *et al.*, 2009, 2015). As a pre-processing step, within each cohort, each trait was quantile normalised, and the effects of age, sex and ten leading principal components of the genetic population structure were regressed out using a linear model. All cohorts were genotyped using Illumina arrays, and imputed by IMPUTE2 (Howie *et al.*, 2009) using the 1000 Genomes Project reference panel (1000 Genomes Project Consortium, 2012). In the analyses, we included 455 521 SNPs on chromosome 1 and, additionally, the SNPs in the following 5 genes:

- *APOE* (apolipoprotein E), 259 SNPs on chr 19;
- *CETP* (cholesteryl ester transfer protein), 387 SNPs on chr 16;
- *GCKR* (glucokinase (hexokinase 4) regulator), 160 SNPs on chr 2;
- *PCSK9* (proprotein convertase subtilisin/kexin type 9), 265 SNPs on chr 1;
- *NOD2* (nucleotide-binding oligomerization domain containing 2), 145 SNPs on chr 16.

We expected that this set of genes would provide a comprehensive spectrum of associations with our phenotypes, since *APOE, CETP, GCKR* and *PCSK9* have well-known associations to lipid levels, whereas *NOD2* is not known to have such an association (NHGRI GWAS catalogue, Hindorff *et al.* (2011)). All SNPs used were of good quality: IMPUTE2 info ≥ 0.8 (Marchini and Howie, 2010) and minor allele frequency ≥ 0.05.

For multi-SNP models, we compared the results from Finnish genotype data with those obtained by estimating the genotypic correlation structure 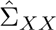 from the 1000 Genomes Project data on 503 European individuals (release 20130502).

For each cohort, genotypic and phenotypic correlation structures computed based on *X*^(*i*)^ and *Y*^(*i*)^, as shown in the eq. (5) and (6), can be found in Supplementary Figures 3-4.

## 3 Results

### 3.1 Performance assessment

The purpose of this section is to validate that *metaCCA* applied to summary statistics produces similar results to the standard CCA (MATLAB function *canoncorr)* applied to the individual-level data.

For *metaCCA*, we always use 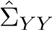 estimated by the method described in section 2.2.2 using summary statistics of the entire chromosome 1.

We focus on the effects of:

- the amount of shrinkage applied to the full covariance matrix *(metaCCA/metaCCA+);*
- estimating 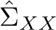from the population underlying the analysis (here, Finnish), or from a more heterogeneous panel (here, European individuals from the 1000 Genomes database).

#### 3.1.1 Univariate genotype — multivariate phenotype

We conducted a meta-analysis of the three cohorts (YFS, FINRISK and NFBC) by testing associations between each SNP in the five genes with different numbers of traits, ranging from 2 to 50. Multi-trait analyses are most useful for correlated traits (Stephens, 2013). To reflect this, for each SNP, we started with a randomly selected trait, and at each step of the analysis, added the trait mostly correlated with the already chosen ones, excluding correlations with absolute values above 0.95. For each SNP, we repeated the procedure three times with different starting lipid measures.

The scatter plot in Figure 2a shows that *metaCCA* applied to the cohort-wise summary statistics provides an excellent agreement with the standard CCA of the pooled individual-level data. Thus, in the one-SNP analysis, we can base the inference on *metaCCA* and put less weight on *metaCCA +* (Figure 2b) that, as expected, produces conservative p-values.

The wide range of the observed −log10 p-values (0 to 88) shows that multivariate association tests can be very powerful in realistic settings, and that our example assesses the performance of *metaCCA* throughout the range that is important in practical analyses. Supplementary Figure 5 further refines the behaviour of *metaCCA* within the range most encountered in genome-wide association studies (0 to 10).

#### 3.1.2 Multivariate genotype — multivariate phenotype

When both genotype and phenotype are multivariate, genotypic correlation structure 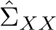needs to be estimated in addition to 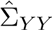. We conducted the meta-analysis of two study cohorts (YFS and NFBC), and computed 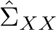 either from FINRISK (FIN) or from 1000 Genomes European individuals (1000G). (Supplementary Table 3 shows errors of 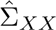 estimates.) We analysed together between 2 to 10 highly correlated lipid measures, chosen sequentially as in the single-SNP tests in section 3.1.1. For each of the five genes, we analysed together between 2 to 10 SNPs that were chosen to be approximately uncorrelated to cover a large proportion of genetic variation within the gene. Each set of SNPs was tested for an association with each group of correlated lipid measures. We repeated the procedure ten times for each gene, with different starting phenotypes and SNPs.

The results are summarised in Figure 2c-f. Figure 2c shows that when genotypic correlation 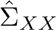 is estimated from the target population, *metaCCA* produces highly consistent results with the standard CCA based on individual-level data. When 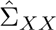 is estimated from a less well matching population (Figure 2e), the accuracy is reduced, and some −log10 p-values become clearly overestimated. In both cases, further shrinkage by *metaCCA +* removes almost completely any overestimation (Figure 2d,f). This property is expected to be important in genome-wide association studies, where *metaCCA+* can protect from false positives when genotypic correlation structure cannot be accurately estimated. *metaCCA+* has less statistical power than the individual-level CCA, but it is still able to detect strong true associations.

Figure 3 illustrates the impact of the number of genotypic and phenotypic features included in the analysis on the accuracy of *metaCCA/metaCCA+.* It shows that further improvement of the agreement with the results of pooled-individual-level analysis is achieved by testing a smaller number of variables jointly.

**Figure 3:**
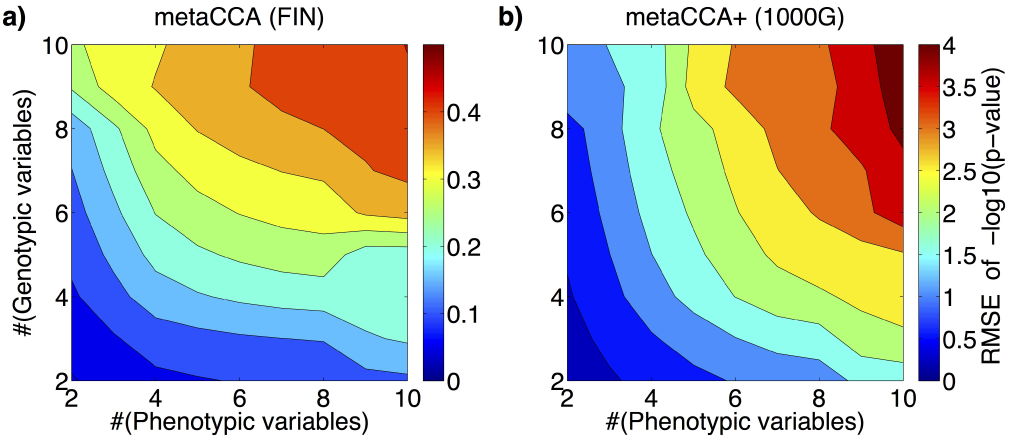
*Multivariate genotype - multivariate phenotype*; Contour plots showing an evolution of root mean square error (RMSE) for a) *metaCCA’s*, b) *metaCCA+’s* −log10 p-values as a function of the number of genotypic and phenotypic features included in the analysis. 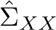 was calculated from a) FINRISK cohort, b) the 1000 Genomes database (EUR individuals), and 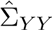 was estimated from summary statistics for SNPs within chromosome 1. Contour plots for up to 25 variables are shown in Supplementary Figure 6.

### 3.2 Application to summary statistics from SNPTEST

In the genetics community, established software packages like SNPTEST (Marchini and Howie, 2010) are used to perform univariate genome-wide tests. In this section, we conduct a meta-analysis of univariate results from standard SNPTEST runs on NFBC and YFS cohorts by *metaCCA.* These cohorts have been meta-analysed previously using individual-level genotypes and the same serum metabolomic profiles that we consider here (Inouye *et al.*, 2012). This single-SNP GWAS highlighted candidate genes for atherosclerosis, and demonstrated the power of incorporating multiple related traits into the analysis. Here, we show that by *metaCCA* we obtain those same results without the access to the individual-level data, and on top of that we can also analyse multiple SNPs jointly by using only summary statistics from the original studies.

We wanted to choose a set of correlated traits for the joint analysis, and therefore we proceeded as follows. By an agglomerative hierarchical clustering (average linkage) of ∑_*Y Y*_ (81 traits) we identified groups of related lipid measures. From the largest of 6 distinct clusters, we selected a set of traits in such a way that no pair exhibited correlation above 0.95. We ended up with a group of 9 lipid measures related to 8 VLDL particles of different sizes and one HDL particle (XXL.VLDL.PL, XXL.VLDL.P, XL.VLDL.PL, L.VLDL.PL, VLDL.D, M.VLDL.FC, S.VLDL.PL, XS.VLDL.TG, S.HDL.TG described in Supplementary Table 1).

We conducted two types of meta-analyses of NFBC and YFS:

1. *Univariate genotype - multivariate phenotype* Each SNP from chromosome 1 tested for an association with the set of 9 correlated lipid measures.
2. *Multivariate genotype - multivariate phenotype* For each of the 5 genes *(APOE, CETP*, *GCKR*, *PCSK9*, *NOD2)*, a group of 5 SNPs, chosen by maximising variation covered with the gene (see Methods 2.4), tested for an association with the set of 9 correlated lipid measures.

The input summary statistics for *metaCCA* were obtained by performing univariate tests for each SNP-trait pair separately using SNPTEST applied to the individual-level data, and transforming the resulting regression coefficients using (4). The correlation structure of analysed traits 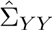 was estimated based on summary statistics for SNPs across the entire genome. The genotypic correlation structure for multi-SNP analyses, 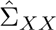, was calculated from the FINRISK cohort.

We compared the results of *metaCCA* and *metaCCA+* with the pooled individual-level CCA of original data sets. Figure 4 shows scatter plots of −log10 p-values for 455 521 SNPs from chromosome 1. The results of *metaCCA* demonstrate an excellent agreement with the original p-values, validating that *metaCCA* can conduct reliable multivariate meta-analysis from standard univariate GWAS software output. As anticipated, *metaCCA+* produces conservative p-values. Manhattan plots illustrating p-values along the chromosome are shown in Supplementary Figure 7. Genome-wide significant associations (at the threshold of *p* = 5 × 10^−8^ standard in the field) are located within two regions: *USP1/DOCK7* and *FCGR2A/3A/2C/3B*, which are known to be associated with lipid metabolism (NHGRI GWAS catalogue, Hindorff *et al.* (2011)). *metaCCA* identified both regions, and *metaCCA+* found the stronger out of the two signals *(DOCK7/USP1).* For top-SNP in *FCGR2A/3A/2C/3B, metaCCA+’s* −log10 p-value is 6.11, compared to 7.73 produced by CCA on individual-level data.

**Figure 4:**
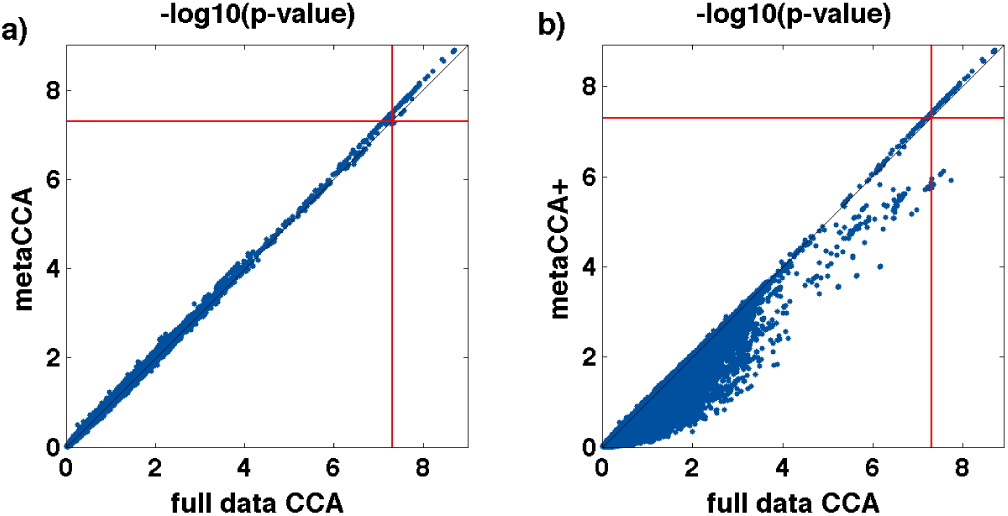
Scatter plots of −log10 p-values from the pooled individual-level CCA of NFBC and YFS and a) *metaCCA*, b) *metaCCA+.* Each point corresponds to one genetic variant from the chromosome 1, tested for an association with the group of 9 correlated lipid measures. In total, 455 521 SNPs were analysed. Red lines indicate the significance level of 5 × 10^−8^ (7.301 on −log10 scale).

Table 1 summarizes the results of the metaanalysis where sets of 5 SNPs, representing each gene, were tested for an association with the group of 9 related lipid measures. Both *metaCCA* and *metaCCA+* produced very accurate p-values (error < 0.2 on −log10 scale) for the four genes with signals below 9. For the largest signal, 9.60 *(CETP)*, −log10 p-values were about one unit overestimated *(metaCCA)* or underestimated *(metaCCA+).* Any of these differences would be unlikely to lead to false inferences when a reference significance level in a gene-based analysis was set to 0.05/20000 = 2.5 × 10^−6^, i.e., 5.61 on −log10 scale, based on there being about 20 000 protein-coding genes in the human genome. At this level, both *metaCCA* and *metaCCA+* found an association between *APOE* and *CETP* and the network of VLDL and HDL particles studied. For *GCKR* and *PCSK9*, the gene-based test did not yield strong signals, even though Table 1 shows that there are individual SNPs in these genes with relatively strong signals. Thus, some other procedure for picking SNPs for a joint analysis might be more fruitful at these genes (see Discussion). An advantage of *metaCCA* framework is its adaptability to any such procedure. Note that *NOD2* has no (known) association with metabolic traits, and therefore it serves as a negative control in Table 1.

**Table 1:** *Multi-SNP:* Comparison between −log10 p-values of CCA on pooled individual-level data sets (NFBC+YFS) and the meta-analyses conducted using *metaCCA* and *metaCCA+.* For each gene, a set of 5 SNPs was tested for an association with the group of 9 related lipid measures. *Top single-SNP:* For each gene, the largest −log10 p-value from single-SNP-multi-trait *(CCA)* and all possible single-SNP-single-trait *(univariate)* tests are shown (with details in Supplementary Table 4). The number of tests in each gene is 1 for *multi-SNP* tests, *G* for *CCA* and 9 × *G* for *univariate*, where *G* is the number of SNPs in that gene.

| gene | multi-SNP |  |  | top single-SNP |  |
| --- | --- | --- | --- | --- | --- |
|  | CCA | *metaCCA* | *metaCCA+* | CCA<br>(multi-trait) | univariate<br>(single-trait) |
| *APOE* | 8.36 | 8.51 | 8.25 | 11.54 | 8.60 |
| *CETP* | 9.60 | 10.60 | 8.51 | 23.77 | 10.04 |
| *GCKR* | 3.95 | 3.89 | 3.80 | 9.64 | 14.21 |
| *PCSK9* | 0.12 | 0.13 | 0.12 | 6.58 | 4.63 |
| *NOD2* | 0.22 | 0.27 | 0.20 | 0.97 | 2.46 |

## 4 Discussion

The advantage of multivariate testing of genetic association is well reported in the literature (Stephens, 2013; Inouye *et al.*, 2012), and also demonstrated in our results (e.g. *CETP* in Table 1 that has multivariate p-value 13 orders of magnitude smaller than any of the univariate p-values). Optimal use of correlated traits is becoming increasingly important as high-throughput phenotyping technologies are being more widely applied to individual study cohorts and large biobanks (Soininen *et al.*, 2015).

We introduced *metaCCA*, a computational approach for the multivariate meta-analysis of GWAS by using univariate summary statistics and a reference database of genetic data. Thus, our framework circumvents the need for complete multivariate individual-level records, and tackles the problem of low sample sizes in individual cohorts by a built-in meta-analysis approach. To our knowledge, *metaCCA* is the first summary statistic-based framework that allows multivariate representation of both genotypic and phenotypic variables.

In large meta-analytic efforts, the ability to work with summary statistics is beneficial, even when there is an access to the individual-level data. For example, with a study design of the Global Lipids Genetics Consortium (2013), we estimate that the reduction in the size of input data between *metaCCA* and standard CCA could be over 750-fold (Supplementary Figure 1).

We provided two variants of the algorithm: *metaCCA* and *metaCCA+.* Based on our results, *metaCCA* is the method of choice when the accuracy of estimated correlation matrices 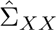 and 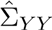 is good, e.g., 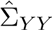 estimated from at least one chromosome, and 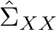 estimated from genetic data from the target population. In such cases, p-values from *metaCCA* were accurate, meaning that false positive and false negative rates are close to those of standard CCA applied to the individual-level data. When the quality of the estimates 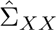 and 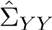 was reduced, *metaCCA+* proved useful to protect from an increase of false positive associations (Figure 2 and Supplementary Figure 8). This is important in GWAS context, where false positives could lead to considerable waste of resources in subsequent experimental and functional studies. A topic for future work would be to further develop our current elbow heuristic of *metaCCA +* to decrease its false negative rate without sacrificing its good false positive rate.

We derived the framework assuming that all traits within each cohort have been measured on the same number of individuals (*N*). While a small proportion of missing data for each trait could be handled by statistical imputation methods, further work is required to study how *metaCCA* should be applied when the sample sizes between the traits vary considerably. We note that the distribution of the test statistic depends on *N* (Supplementary Data), as do the effect sizes transformation (eq. 4) and meta-analysis approach (section 2.3).

For multivariate phenotype data, several types of association tests are possible. Natural question is which one should we prefer in practice. It is evident that single-SNP-multi-trait tests can detect much stronger signals at some SNPs than any of the univariate tests separately (e.g. *CETP* in Table 1), and identify associations not found by univariate approach (Inouye *et al.*, 2012). On the other hand, for some other SNPs, the highest univariate signal may be clearly higher than the multi-trait one, even after accounting for the increase in the number of tests. For example, in *GCKR* (Table 1), the top SNP’s (rs1260326) association was explained already by one of the traits individually (M.VLDL.FC). Given the difference in degrees of freedom of the tests, this led to a 4.6 units higher −log10 p-value in the univariate test compared to the multivariate one. Thus, for single-SNP tests, univariate and multivariate tests complement each other and neither should be excluded from consideration.

When also genotypes are multivariate, even more possibilities for association testing emerge. Multigenotype tests are common practice in rare variant association studies, where statistical power to detect any single variant is very small (Lee *et al.*, 2014; Feng *et al.*, 2014). The set of variants is often chosen based on functional annotation, such as predicted nonsense or missense effects. In this paper, we have rather focused on common variants (MAF > 5%) to ensure the accuracy of the summary statistics and genotypic correlations used in *metaCCA.* To illustrate our multi-SNP approach, we chose a fixed number of SNPs that tag a maximal amount of genetic variation within a gene. However, *metaCCA* could equally well incorporate any other way of choosing the SNPs, for example, motivated by functional annotation (ENCODE Project Consortium, 2012), known expression effects (Ardlie *et al.*, 2015) or previous GWAS results on other traits (Hindorff *et al.*, 2011). A topic for further research could be to extend the covariance matrix-based analyses from CCA to dynamic approaches that learned from the data the set of variants and traits to be considered together. This would circumvent the need to restrict the subset of variables before the analysis.

We envision that multivariate association testing using *metaCCA* has a great potential to provide novel insights from already published summary statistics of large GWAS meta-analyses on multivariate high-throughput phenotypes, such as metabolomics and transcriptomics. Finally, we hope that our work helps extending the application area of CCA to summary-statistic data also in other data-rich fields outside genetics.

## Acknowledgement

This work was financially supported by the Helsinki Doctoral Education Network in Information and Communications Technology (HICT) to AC, the Academy of Finland [257654 to MP; 251170 to the Finnish Centre of Excellence in Computational Inference Research COIN; 259272 to PM]. SR was supported by the Academy of Finland (251217 and 255847), Center of Excellence in Complex Disease Genetics, EU FP7 projects ENGAGE (201413) and BioSHaRE (261433), the Finnish Foundation for Cardiovascular Research, Biocentrum Helsinki, and the Sigrid Juselius Foundation.

